# A ghost of dioecy past and the legacy of sexual dimorphism: low siring success of hermaphrodites after the breakdown of dioecy

**DOI:** 10.1101/430041

**Authors:** Luis Santos del Blanco, Eleri Tudor, John R. Pannell

## Abstract

Evolutionary transitions from dioecy to hermaphroditism must overcome the inertia of sexual dimorphism because modified males or females will express the opposite sexual function for which their phenotypes have been optimized. We tested this prediction by comparing the siring success of female-derived hermaphrodites of the plant *Mercurialis annua* with males and hermaphrodites that present a male-like inflorescence. We found that pollen dispersed by female-derived hermaphrodites was about a third poorer at siring outcross offspring than that from hermaphrodites with male-like inflorescences, illustrating the notion that a ‘ghost of dioecy past’ compromises the fitness of derived hermaphrodites in outcrossing populations. We conclude that whereas dioecy might evolve from hermaphroditism by conferring upon individuals certain benefits of sexual specialization, reversals from dioecy to hermaphroditism must often be limited to situations in which outcrossing cannot be maintained and inbreeding is favored. Our study provides novel empirical support for evolutionary models for the breakdown of dioecy.

## Introduction

Dioecy is found in about 6% of species, but in about half of all families of flowering plants (Renner & Ricklefs 1995; Weiblen *et al*. 2000; Renner 2014). This distribution might suggest that dioecy is an evolutionary dead end or ‘failure’ (Westergaard 1958; Bull & Charnov 1985; Heilbuth 2000), such that lineages with separate sexes diversify less and are more prone to extinction than their hermaphroditic counterparts. However, the dead-end hypothesis has been challenged by analysis suggesting that the evolution of dioecy might actually increase lineage diversification, and that the scattered phylogenetic distribution of dioecious species might be explained by frequent reversions from dioecy to functional hermaphroditism, i.e., to a state in which plants have either bisexual flowers or flowers of both sexes (‘monoecy’) (Kafer *et al*. 2014; Käfer & Mousset 2014; Sabath *et al*. 2016; Kafer *et al*. 2017).

Evidence for reversions from dioecy to hermaphroditism has been accumulating for some time. Phylogenetic analysis of clades in which both dioecy and hermaphroditism occur indicates that dioecy was probably not derived from, but was ancestral to, hermaphroditism (Goldberg *et al*. 2017; Kafer *et al*. 2017), including, e.g., the large and successful Cucurbitaceae family, in which it was lost several times (Volz & Renner 2008; Schaefer & Renner 2010), and in the genera *Bursera* (Becerra & Venable 1999), *Garcinia* (Sweeney 2008), *Gallium* (Soza & Olmstead 2010) and *Dodonaea* (Harrington & Gadek 2010). Within genera, Goldberg et al. (2017) found that transitions towards combined sexes were no less common than those towards separate sexes. In many of these reversions, dioecy evolved from monoecy, not hermaphroditism with bisexual flowers, suggesting that the association between dioecy and monoecy (Renner & Ricklefs 1995) may be explained not only by the evolution of dioecy from hermaphroditism via monoecy, but also by the breakdown of dioecy to monoecy.

The breakdown of dioecy presumably involves the selection of individuals with ‘leaky’ sex expression, i.e., males producing occasional fruits and seeds, or females producing occasional flowers with functional stamens and pollen. Leaky sex expression is common in dioecious populations (reviewed in Ehlers & Bataillon 2007; Cossard *et al*. 2018), and has been invoked in models for the breakdown of dioecy in both plants (Crossman & Charlesworth 2014) and animals (Pannell 2008). Lloyd (1975a) suggested that dioecy had broken down in *Leptinella* as a result of selection of leaky males following colonization. Similarly, monoecy in the *Mercurialis annua* species complex probably evolved from ancestral dioecy (Krahenbuhl *et al*. 2002; Obbard *et al*. 2006) under selection for reproductive assurance in metapopulations with frequent colonization (Pannell 2001).

Although the breakdown of dioecy clearly involves changes in sex expression and sex allocation, the relative fitness of males, females and invading hermaphrodites must also depend on the extent to which the dioecious population is sexually dimorphic for non-reproductive traits. Sexual dimorphism is almost ubiquitous in dioecious plants, with males and females differing in morphological, physiological, defense, life-history, resource acquisition and inflorescence traits (Darwin 1871; Geber *et al*. 1999; Fairbairn *et al*. 2007; Moore & Pannell 2011; Barrett & Hough 2013). The evolution of sexual dimorphism likely allows males and females in dioecious populations to express phenotypes that enhance siring success and seed production, respectively, but that might compromise these functions if expressed in the other sex (Lande 1980; Cox & Calsbeek 2009). Yet reversion to hermaphroditism must bring about just this sort of compromise, because modified males or females will express their newly acquired function in the context of a phenotype optimized for the opposite sex. The evolution of sexual dimorphism should constrain the breakdown of dioecy in a way that goes beyond a simple sex-allocation trade-off.

The likely constraints of sexual dimorphism on the breakdown of dioecy are well illustrated in wind-pollinated species, in which male and female inflorescences are often quite different (Lloyd & Webb 1977; Whitehead 1983; Weberling 1992; Galonka *et al*. 2005; Friedman & Barrett 2009b; Harris & Pannell 2010; Harder & Prusinkiewicz 2013). Male (or staminate) flowers of wind-pollinated species are typically held on flexible stalks or ‘peduncles’, which facilitate pollen liberation from anthers and pollen dispersal by wind (reviewed in Harder & Prusinkiewicz 2013). In trees, these structures often hang from the branches. In herbs, they are typically held above the plant canopy. In both situations, pollen is liberated when turbulent gusts shake male flowers or anthers (Urzay *et al*. 2009). In contrast, female (or pistillate) flowers are typically held on more rigid inflorescences, and pollen is picked up by stigmas as they impact its surface, or from non-viscous eddies around the flower (but see Cresswell *et al*. 2010). Whereas the two sexual functions of bisexual hermaphrodite flowers will often interfere with one another in wind-pollinated plants, compromising fitness (Friedman & Barrett 2009b; Harder & Prusinkiewicz 2013), the inflorescences of plants with separate male or female flowers (in dioecious or monoecious species) may be optimized for each sex separately (Friedman & Barrett 2008), e.g., male flowers of *Zea mays* are born on flexible tassels at the shoot apex, whereas female flowers develop in the leaf axils (Aylor *et al*. 2003).

The male and female inflorescences of the plant *Mercurialis annua* L. (Euphorbiaceae) illustrate the divergent strategies for the two sexes of wind-pollinated herbs and suggest how they might constrain the breakdown of dioecy (Figure 1a). *Mercurialis annua* is a complex of European ruderal plants that vary in their sexual systems and inflorescences (Durand 1963; Durand & Durand 1992; Pannell *et al*. 2008). Dioecy is ancestral in *Mercurialis* and is widespread in *M. annua* in Europe, but monoecy has apparently evolved from dioecy in the Iberian Peninsula and north Africa through the spread of leaky females with an enhanced male function (Obbard *et al*. 2006). In dioecious populations, males disperse their pollen from staminate flowers on ‘pedunculate’ stalks held above the foliage, whereas females produce their flowers on subsessile pedicels in leaf axils (Eppley & Pannell 2007). The male function of the monoecious *M. annua* is associated with a female phenotype that differs from males in terms of life-history, nitrogen budget and allocation to defense (Hesse & Pannell 2011b, c; Sanchez-Vilas & Pannell 2011a; Sanchez-Vilas & Pannell 2011b; Sanchez-Vilas *et al*. 2011; Labouche & Pannell 2016; Tonnabel *et al*. 2017). Importantly, monoecious individuals hold both their male and female flowers in leaf axils, whereas males place flowers on erect inflorescence stalks (‘peduncles’) (Figure 1a). Pollen dispersed from male peduncles is a 60% better at siring outcrossed progeny than that from monoecious inflorescences (Eppley & Pannell 2007). We might view the poor pollen dispersal of monoecious individuals as a ‘ghost of dioecy past’, in the sense that they are ‘haunted’ by the legacy of sexual dimorphism inherited from their dioecious ancestors.

**Figure 1.**
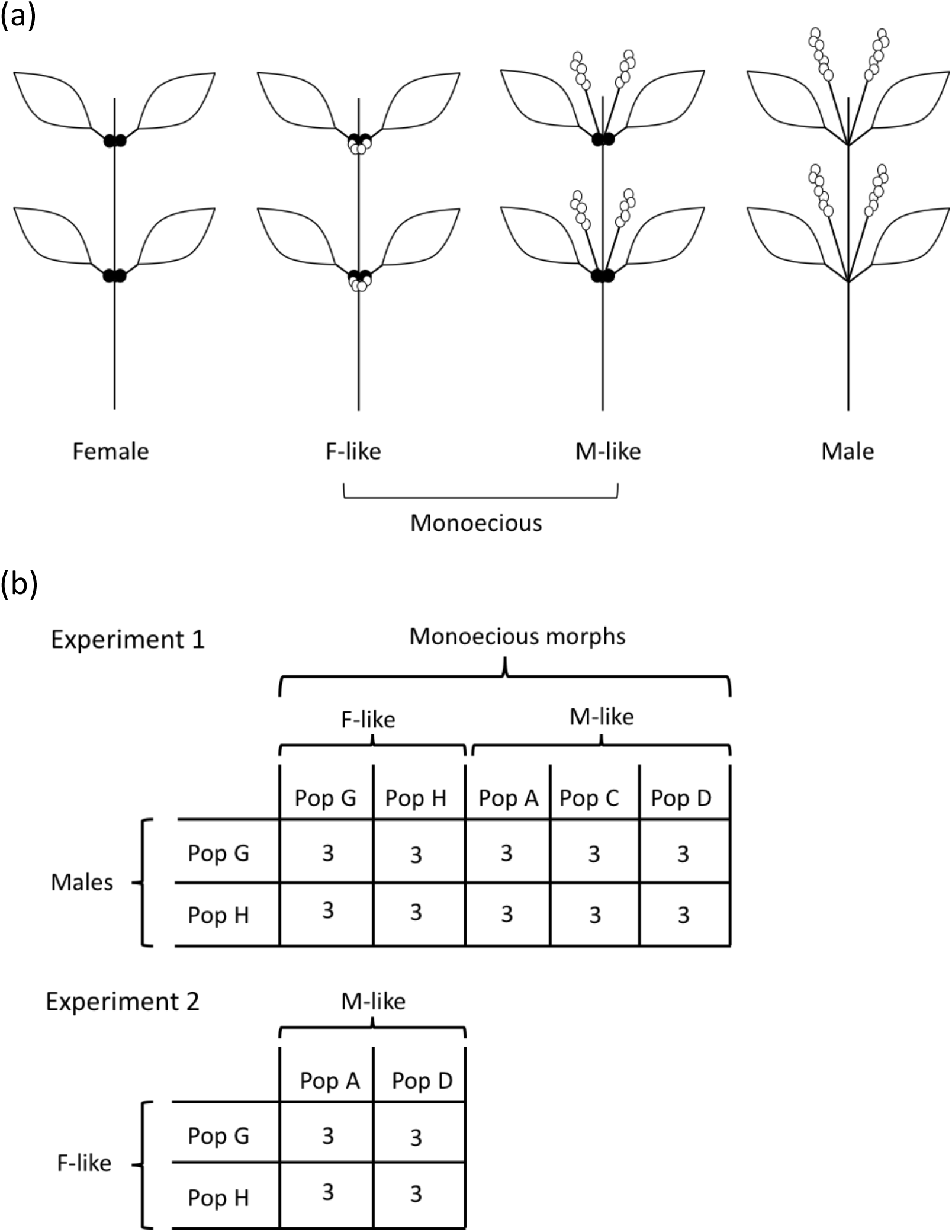
(a) Cartoon drawings illustrating the inflorescence variation in *Mercurialis annua* between females, F-like monoecious individuals, M-like monoecious individuals, and males. Black circles represent female flowers or fruits. White circles represent male flowers. (b) Experimental design for Experiments 1 and 2. There were three replicates for each combination of competing populations indicated. In Experiment 1, males from each of two androdioecious populations competed either against female-like monoecious individuals from each of the same two populations, or male-like monoecious individuals from each of three populations. In Experiment 2, female-like monoecious individuals from each of two of their populations competed against male-like monoecious individuals from each of two of their populations.

Here, we compare the siring success of the typical monoecious individuals with female-like (‘F-like’) inflorescences with that of hitherto undescribed monoecious individuals of *M. annua* that have longer inflorescences similar to those of males (‘M-like’). Populations of the F-like form are widespread around the coast of the Iberian Peninsula, whereas populations of the M-like form are restricted to southern and eastern Spain. Although the two forms rarely co-occur, their distributions are broadly sympatric. The evolutionary paths linking dioecy to the two monoecious forms in *M. annua* are not well understood. However, pedunculate inflorescences are associated with a Y-linked marker in all lineages that have them, except in M-like monoecious individuals (unpublished data). It is thus likely that peduncles of the M-like lineage are not derived directly from males and have evolved independently. Either way, the peduncle represents a potential improvement on the female-derived monoecious inflorescence.

We hypothesized that the M-like form should enjoy greater siring success than the F-like form, in competition both with males with the F-like form. We first compared inflorescence morphology and pollen production of males with M-like and F-like monoecious forms, then tested our hypotheses by evaluating siring success of the three phenotypes in common gardens. We also tested for trade-offs between male and female allocation within and among populations. Our results support the notion that monoecious individuals that retain a female-derived morphology are indeed poorer at siring progeny than those that combine monoecy with male-like inflorescences, illustrating a ghost of dioecy past. Our analysis also suggests that transitions from dioecy to hermaphroditism are likely to be associated with a shift from obligate outbreeding to facultative inbreeding, as implied by models of the breakdown of dioecy (Maurice & Fleming 1995; Wolf & Takebayashi 2004; Ehlers & Bataillon 2007; Pannell 2008; Crossman & Charlesworth 2014).

## Materials and Methods

### Study populations, seed collection, and seedling establishment

We collected seed from about 30 seed-producing individuals from each of five populations: two androdioecious populations in which males co-occurred at frequencies of approximately 30% with F-like monoecious individuals (populations G and H); and three monoecious populations with only M-like monoecious individuals (populations A, C and D). We sowed seeds in bulk in seedling trays and transplanted seedlings (7 – 10 days after sowing) individually into small pots. When their sex could be determined (at 21 days), we re-transplanted them into 15 cm diameter pots and established them in their mating arrays outside.

### Experimental design

We conducted two experiments to compare male and female components of fitness: in Experiment 1 (conducted at Wytham field station, University of Oxford), each monoecious form competed separately against males; in Experiment 2 (conducted at the University of Lausanne), we competed the two monoecious forms against one another (Figure 1b). For Experiment 1, we placed males from each androdioecious population with individuals of either the M-like and F-like monoecious forms, yielding ten different common-garden combinations. Each combination was replicated three times (30 mating arrays in total), with seven males and 42 monoecious individuals arranged so that males occurred only once in each row and column. Plants were allowed to mate with one another for four to six weeks, when they were harvested, with seeds collected from each mother separately. (Note that any seeds sired prior to establishment of the respective arrays had already dispersed, so that we can be sure that all progeny harvested were sired under the treatment conditions.)

In Experiment 2, mating arrays comprised individuals of both monoecious forms together. We used genotypes from two F-like (G and H) and M-like populations (A and D), competing in all four combinations (A-G, A-H, D-G and D-H), with three replicates each (twelve arrays in total). Arrays were established as squares with 25 F-like alternating with 24 M-like individuals. Plants were harvested after six weeks.

In both experiments, arrays were established across the available area. Arrays for Experiment 1 were tens of meters apart, but for Experiment 2 they had to be placed at approximately three meters from each other. To prevent immigration of pollen from adjacent arrays, we erected plastic barriers 1.5 m tall between them. A similar setup prevented gene flow among contiguous plots in a previous experiment (Dorken & Pannell 2009). Any gene flow among arrays would have reduced experimental power, so our results are conservative.

### Data collection

Individuals in the outer edge of all arrays, except males in Experiment 1, were excluded from analysis. Experiment 1 had a final sample size of 851 plants (278 F-like, 373 M-like, and 210 male individuals); Experiment 2 had a final sample size of 300 plants (156 F-like and 144 M-like individuals). We measured the height of all plants, disregarding the additional height of pedunculate inflorescences. For a subsample of seven plants of each morph and array, we measured the length of five randomly chosen inflorescences (both experiments), and the biomass of all staminate flowers. Previous work has shown that pollen biomass is strongly correlated with the biomass of staminate flowers (Pannell 1997b, c). All plants from both experiments were allowed to dry slowly and release their seeds porous bags. We weighed the seeds of each plant together, as well as the aboveground plant biomass.

### Assessment of relative siring success

For Experiment 1, we used the sex ratio in the progeny to estimate the relative siring success of males and monoecious individuals. In *M. annua*, males are determined by the expression of a dominant allele at a single locus, so that males are heterozygous (i.e., XY) (Russell & Pannell 2015; Veltsos *et al*. 2018). Progeny sired by males will thus be 50% male and 50% monoecious. Accordingly, we calculated the relative siring success of males in each array as twice the frequency of male progeny, based on 200 progenies grown to sexual maturity per array (6,000 individuals in total).

For Experiment 2, we used microsatellites to estimate selfing rates and siring success by F-like- and M-like individuals. We genotyped 30 plants from progeny produced by each of the two monoecious forms from each of the twelve replicate arrays (total of 720 plants), using DNA from young silaca-dried leaves. DNA was extracted with a BioSprint 96 robot (Qiagen), using a BioSprint 96 DNA plant kit (Qiagen). Individuals were scored for five microsatellite markers (Mh14, Mh15, Mh19, Mh52 and Mh91(2) that provide good separation between the monoecious populations sampled (Korbecka *et al*. 2010). All five markers were amplified in a single multiplexed PCR reaction, following the protocol described in Korbecka et al. (2010). We processed the samples in an ABI 3100 sequencer (Applied Biosystems), and analysed the results with GeneMapper v.4.0 (Applied Biosystems). Individual genotypes were classified as having been sired by a father of the same or a competing phenotype in the array; resolution was insufficient to assign paternity to specific individuals. For populations of the F-like form, we estimated selfing rates using the software RMES (David *et al*. 2007); those of the M-like form had almost no genetic variability.

### Data analysis

We used mixed models to analyse: plant height; plant biomass; mean length of peduncles per plant; pollen biomass and seed biomass; sex ratio in the progeny of Experiment 1; and, for Experiment 2, the proportion of progeny of parents with different phenotypes. As our primary interest was to determine the functional effect of two contrasting monoecious inflorescence forms in *M. annua*, we defined inflorescence form as a fixed variable, and population within inflorescence form and array as random variables. We tested for a trade-off between male and female reproduction of monoecious individuals in Experiment 1 by fitting a model with pollen mass as the response variable and biomass of seeds, plant biomass and inflorescence form as independent variables, controlling for population and array variation as random terms.

We used Gaussian models for all variables except for the sex ratio and inter-form crossing rate, for which we used a binomial model. Data were log-transformed when necessary to normalize residuals. All data analysis was implemented in R v.3.3.2 (R Development Core Team 2016) using package lme4 (Bates *et al*. 2015). Significance of fixed effects and differences between inflorescence forms were evaluated using the package lmerTest (Kuznetsova *et al*. 2017), or z-tests in the case of ratios. Significance of random effects was assessed by likelihood-ratio tests.

## Results

### Phenotypic variation among inflorescence forms

Phenotypic measurements of the three phenotypes (males, F-like- and M-like individuals) were broadly consistent across both Experiments 1 and 2; for brevity, we therefore report measurements for Experiment 1 in the text (Figure 2) and present all data for both experiments in the Supplementary Materials (Tables S1 and S2).

**Figure 2.**
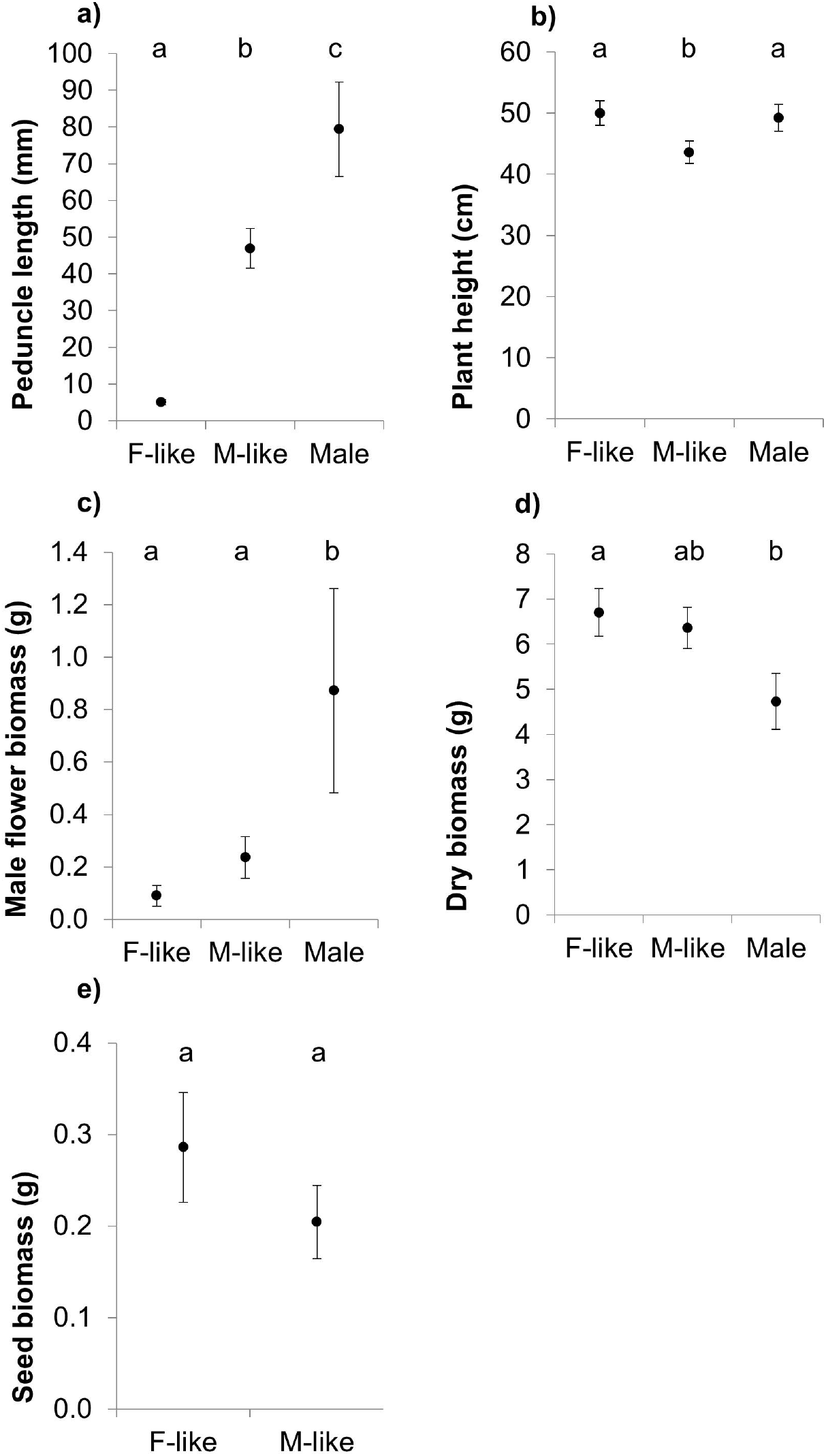
Mean values for several size and allocation traits for males, female-like (F-like) and male-like individuals (M-like) of Mercurialis annua: (a) height including inflorescence; (b) dry biomass; (c) male flower biomass; (d) total biomass of seeds produced; (e) peduncle length. Measurements were taken from mating arrays in Experiment 1 under uniform environmental conditions. Error bars show the standard error. Significant pairwise differences (p < 0.05) are indicated by different low case letters. Sample sizes for panels (b), (d) and (e): males = 210, F-like = 278, M-like = 373. Sample sizes for panels (a) and (c): males = 210, F-like = 84, M-like = 126.

Peduncle length differed between all three phenotypes: peduncles of males were longer than those of M-like monoecious individuals, which were much longer than those of F-like individuals (Figure 2a and Table S1). F-like monoecious individuals were similar in height to males, but significantly taller than M-like monoecious individuals (Figure 2b and Table S1). Males invested much more in male flower production than did both monoecious forms (Figure 2c). Even though there were substantial differences in both vegetative and reproductive traits at the population level (biomass: χ^2^_1_ = 30.6, p < 0.001; seed biomass: χ^2^_1_ = 3.47, p = 0.062; male flower biomass: χ^2^_1_ = 51.2, p < 0.001; Tables S1 and S2), large within-form population variation and low population number rendered differences among forms non-significant (biomass: t1.06 = 0.50, Figure 2d; p = 0.30; seed biomass: t_16_ = 1.40, Figure 2e; p = 0.18; pollen biomass: t_4_ = 1.99, p = 0.12, Figure 2c).

### Inferred siring success

In Experiment 1, in which one or other of the two monoecious forms co-occurred with males, there were substantially more monoecious progeny (as opposed to males) from arrays with M-like (proportion = 0.92+-0.02) than with F-like individuals (proportion = 0.68 +- ± 0.05; z = 6.22, p < 0.001; Figure 3a). Given that maleness in *M. annua* is determined by a dominant allele (see Materials and Methods), we thus inferred that M-like individuals sired 35% more progeny when competing with males than F-like individuals did. Equivalently, F-like individuals sired 25% fewer progeny than their F-like counterparts.

**Figure 3.**
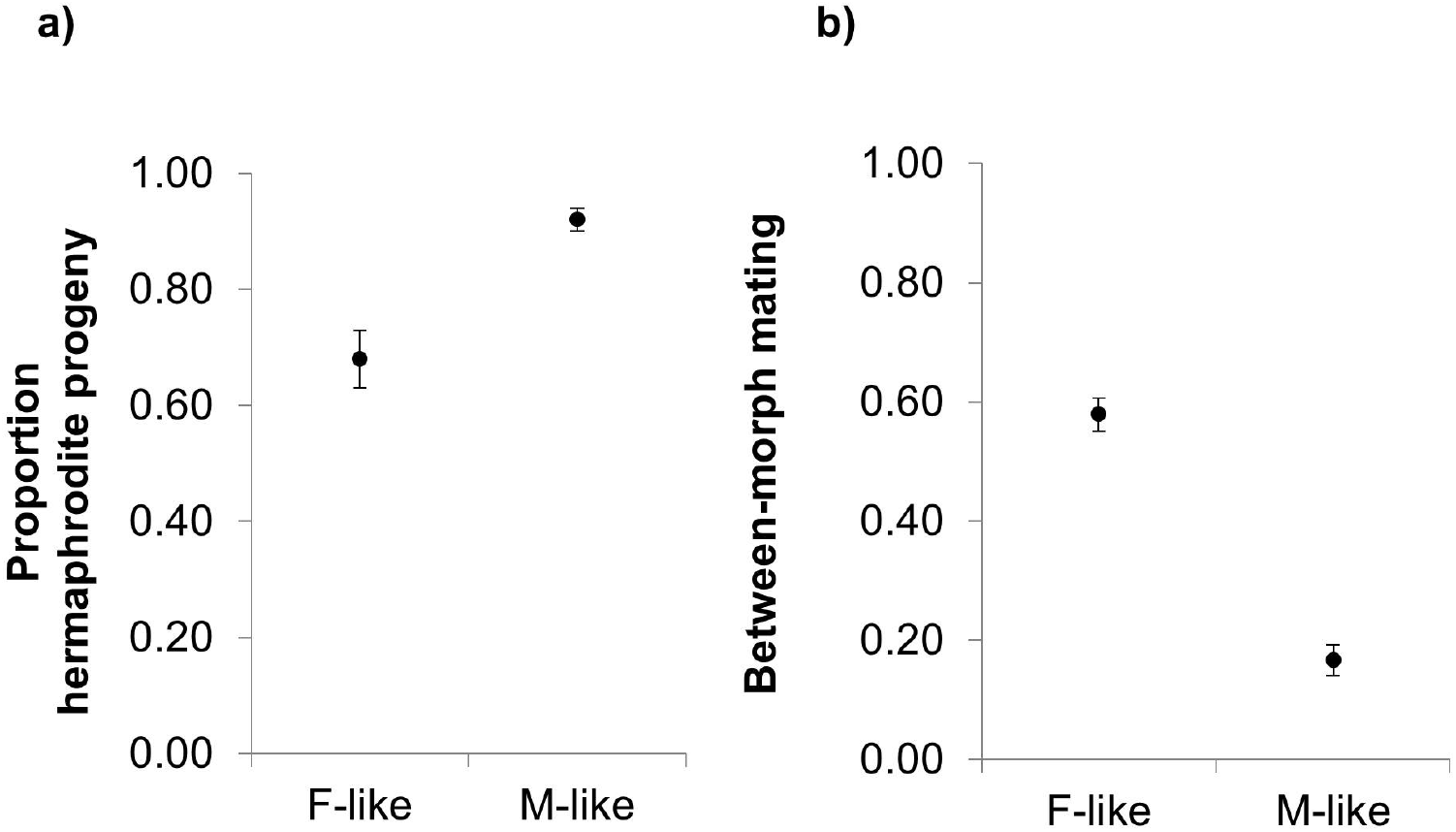
(a) Mean proportion of males in the progeny of female-like (F-like) and male-like (M-like) hermaphrodites grown in mating arrays together with males (Experiment 1). (b) Proportion of progeny of female-like and male-like individuals attributable to crosses with the other morph after mating in mixed mating arrays in Experiment 2. Significant pairwise differences (p < 0.05) are indicated by different low case letters. Samples sizes: (a) M-like = 3600, F-like = 2400; (b) M-like = 360, F-like = 360.

In Experiment 2, in which individuals of the two monoecious forms co-occurred, M-like individuals sired more than three times greater than F-like individuals did (z = 9.3, p < 0.001; Figure 3b). Of progeny with F-like individuals as both parents, 35% were self-fertilized. Taking into account progeny of crosses between the two forms, F-like mothers thus self-fertilized a fraction 0.15 of their progeny (Table S3). There was insufficient variation at the microsatellite loci to estimate the selfing rate of M-like individuals. However, if we assume a similar selfing rate for both forms (see Discussion), we may infer that pollen dispersed by F-like individuals in Experiment 2 sired 31% fewer outcrossed progeny than pollen dispersed by M-like individuals.

### Trade-off between male and female reproduction

There was a significant trade-off between male and female allocation to reproduction within populations (t_4.3_ = 3.47, p = 0.023; Figure 4), with significant variation in the strength of the trade-off among populations (χ^2^_3_ = 22.2, p < 0.001; Figure 4). Biomass (t_184_ = 7.04, p < 0.001) and M-like inflorescence type (t10.1 = 2.39, p = 0.038) had a positive and significant contribution to the model, i.e., individuals with higher biomass and peduncles allocated more to male function for a given female allocation.

**Figure 4.**
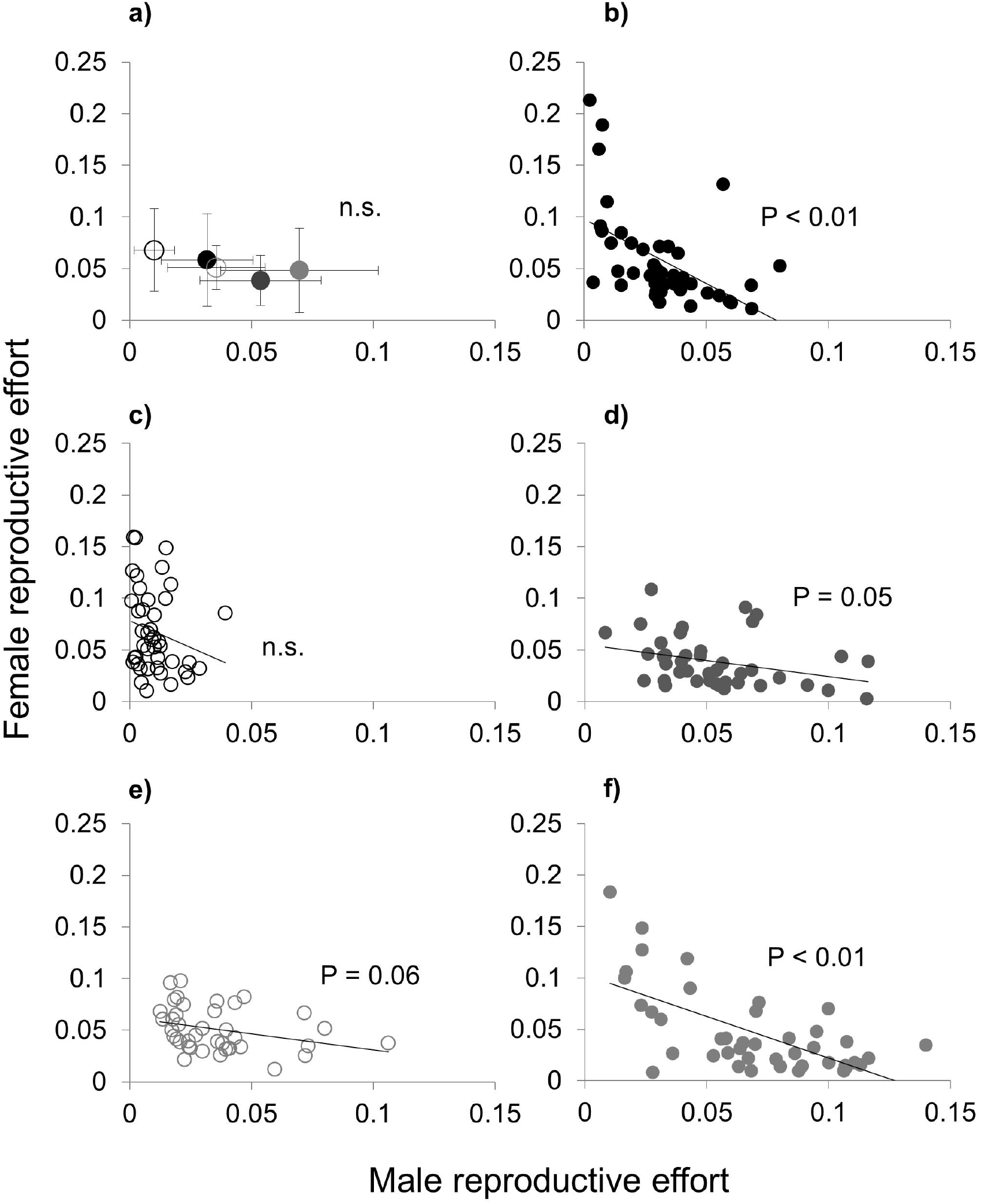
Trade-off between male and female allocation at the individual plant and population level from Experiment 1. (a) Male vs. female average relative reproductive allocation at the population level. Hollow circles represent female-like populations; solid circles represent male-like populations. (b), (d) and (f) male and female relative reproductive allocation at the individual level in each of three populations with male-like individuals (populations A, C and D, respectively). (c) and (e) male and female relative reproductive allocation at the individual level in two female-like populations (populations G and H, respectively). Relative reproductive allocation was estimated as the biomass allocated to either sex divided by total plant biomass.

## Discussion

### High siring success of M-like monoecious individuals

Monoecy in *M. annua* has evolved from dioecy via the modification of females that produce male flowers and that disperse pollen from sub-sessile axillary inflorescences similar to those of females. Our results support the hypothesis that this female-derived inflorescence morphology, a legacy of the breakdown of dioecy, compromises the siring success of monoecious individuals. Monoecious individuals with short female-derived (F-like) inflorescences sired about 25% fewer progeny in mating arrays with males than a newly characterised form with long, male-like (M-like) inflorescences that occurs in part of the species’ range, and they sired 31% fewer progeny when in direct competition with the M-like form.

Our results are coherent with those of Eppley and Pannell (2007), who found that pedunculate inflorescences conferred a 60% siring advantage per pollen grain on males compared with F-like individuals lacking peduncles. Our finding that pollen of the M-like monoecious form has a lower (45%) siring advantage than the F-like form is consistent with both their shorter peduncles than those of males, and their tendency to produce more pollen and fewer seeds than F-like monoecious individuals. The trade-off between male and female allocations within our arrays indicates that individuals in populations emphasising their male function should have a reduced female function.

Our siring estimates assumed that M-like and F-like forms have the same selfing rate. Individuals of the F-like form self-fertilized a proportion 0.15 of their seeds. This relatively low value is consistent with the low selfing rates of F-like monoecious *M. annua* in both dense experimental and field populations (Eppley & Pannell 2007; Korbecka *et al*. 2011). It is also consistent with the presence of males in androdioecious populations of *M. annua*, because males cannot be maintained with hermaphrodites or monoecious individuals that self-fertilize a large proportion of their progeny (Lloyd 1975b; Charlesworth & Charlesworth 1978; Charlesworth 1984). M-like individuals should be less likely to fertilize their own ovules than F-like individuals, so the selfing rate of the former might be that of the latter. M-like individuals produced more pollen than F-like individuals, and selfing rates individuals in *M. annua* correlated positively with pollen production in general (unpublished data). Given these opposing effects of the reproductive strategy of M-like individuals, an assumption of similar selfing rates for the two forms seems reasonable.

The higher sex allocation and siring success of M-like monoecious individuals of *M. annua* helps to explain why they rarely co-occur with males (c.f., Pannell *et al*. 2014), as well as why the two monoecious forms rarely co-occur. Although the weak trade-off among monoecious individuals between their male and female functions might allow co-existence of the two forms (with one emphasising male function and the other emphasising female function), the high fitness of M-like individuals means that F-like individuals should be maintained at low frequencies and could more easily be lost by drift, especially because population size fluctuates so much (Dorken *et al*. 2017). Similarly, although males produce more pollen and have longer peduncles than M-like individuals, our Experiment 1 showed that the siring success of the latter is sufficient to keep males at low frequency; they, too, might thus easily be lost by drift. The effect of drift and demographic stochasticity has similarly been shown to allow the loss of style morphs of *Eichhornia paniculata* that are maintained by negative frequency-dependent selection in large populations (Barrett *et al*. 1989).

The high siring success of the M-like monoecious form of *M. annua* also raises the question of why it has not spread more widely across the Iberian Peninsula, replacing the F-like form. It is possible that the superior mating strategy of the M-like form is costly in ways we have not evaluated, e.g., in terms of physiological and life-history traits that are sexually dimorphic in dioecious or androdioecious *M. annua* (Hesse & Pannell 2011c; Sanchez-Vilas & Pannell 2011b; Sanchez-Vilas *et al*. 2011; Labouche & Pannell 2016; Tonnabel *et al*. 2017). For example, the F-like form might enjoy an advantage over the M-like form during periods of colonization (e.g., because it confers greater reproductive assurance, Friedman & Barrett 2009a). Such an explanation would be consistent with the metapopulation model proposed by Pannell (1997a), notably if greater seed production by the F-like form allowed it to establish more viable demes early after colonisation (Pannell 2001). Estimates of relative seed production and progeny performance from a wider sample of populations would help to evaluate this possibility.

### Sexual dimorphism and the stabilization of dioecy

Variation in inflorescence morphology in *M. annua* illustrates how sexual dimorphism might stabilize the maintenance of separate sexes in dioecious plants. Sex-allocation theory predicts that if either the male or the female sexual function (or both functions together) have accelerating fitness gain curves, then the ‘fitness set’ is concave, and dioecy should be evolutionarily stable (Charnov *et al*. 1976; Charnov 1982; West 2009). Sexual dimorphism likely allows unisexuals to perform better than hermaphrodites in their corresponding sexual function, which should enhance the concavity of the fitness set (Figure 5). Due to lack of specialised structures for pollen dispersal, the male fitness gain curve of F-like hermaphrodites in *M. annua* is probably saturating. In contrast, our results suggest that the superior inflorescence structure of the M-like morph likely diminish this saturating tendency. (Saturation of female fitness gain curves likely remain unchanged.) If so, we suggest that the fitness set of the M-like form should relax conditions for their invasion into a dioecious population and a transition to functional hermaphroditism, particularly under intense competition for outcross siring (Pannell 2001).

**Figure 5.**
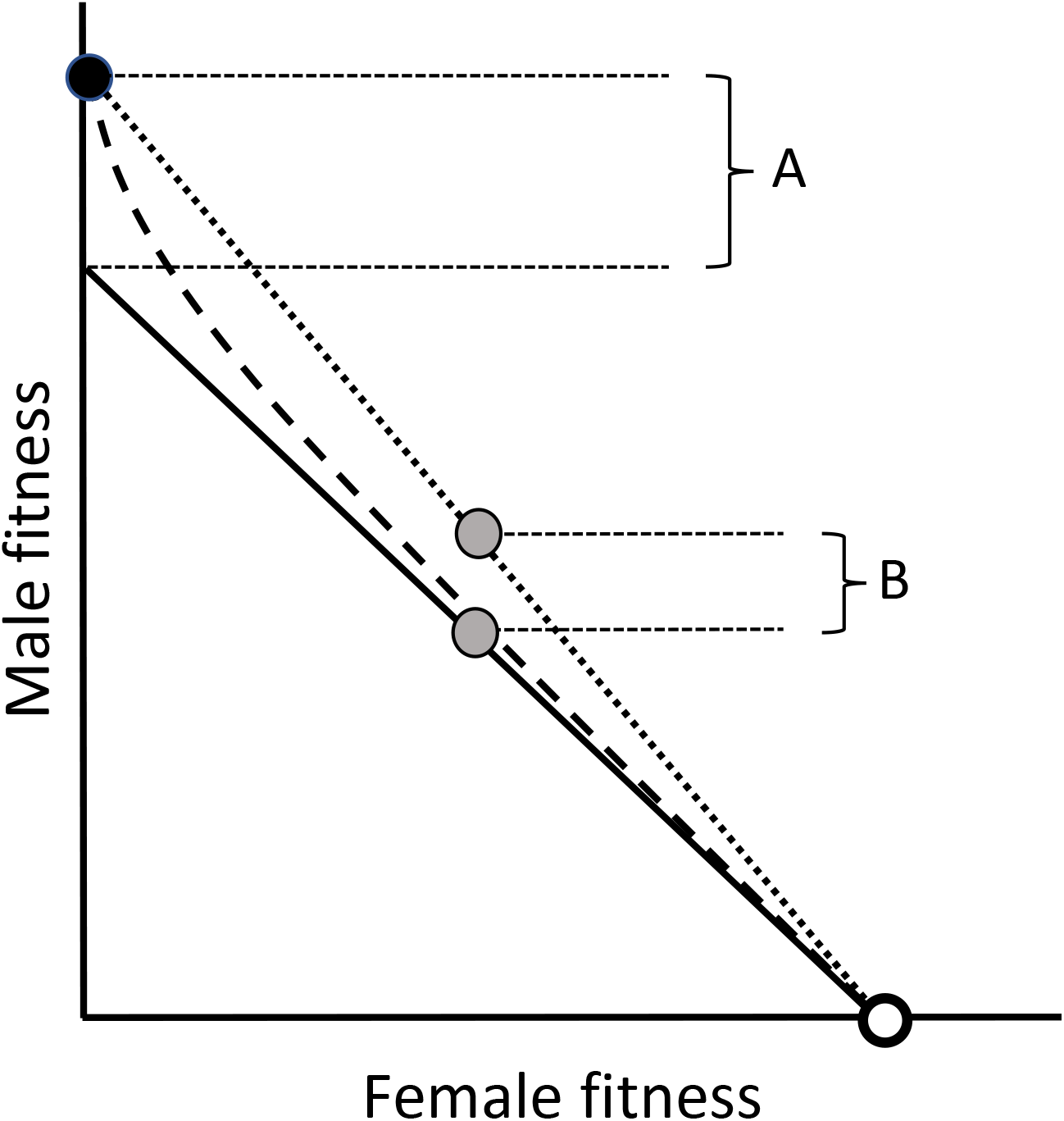
Heuristic schema illustrating the hypothesized influence of the adoption of a specialized male-like inflorescence by males or hermaphrodites on the fitness set relating male to female components of fitness. The diagonal straight line depicts a linear fitness set representing a simple tradeoff between male and female functions of a wind-pollinated plant like *Mercurialis annua*, in the absence of any sexual specialization by males or females. Females (open white circle) occupy the extreme female end of this line. Female-like individuals allocating resources to pollen would occupy positions along the straight-line diagonal. Adoption of a specialized male inflorescence increases the male component of fitness (closed black circle), so that males have a fitness greater than that achievable simply by allocating all reproductive resources to male function (represented by the interval A). The adoption by males of the specialized inflorescence effectively renders the fitness set concave (dashed curve). Adoption by hermaphrodites of a specialized male-like inflorescence increases their male fitness component above that achievable by female-like hermaphrodites (interval B), effectively removing the concavity of the fitness set (dotted line). In our experiments, intervals A and B increased male fitness by about 60% and 45% above the straight-line diagonal. See text for details.

Figure 6 sets out a path that might commonly be followed when dioecy breaks down. In recently evolved dioecious populations, the fitness of males and females will be compromised by the expression of genes in individuals of one sex that are better suited to performance of the other, or genes underpinning a reproductive, physiological, life-history, or defence strategy that is not optimised for its own sex. Over time, selection should act to reduce trait correlations between males and females (Lande 1980), optimising male and female phenotypes differently in a sexually dimorphic population (Figure 6a). It is also possible that selection for sexual specialisation sometimes coincides with the transition from hermaphroditism to dioecy (Willson 1979; Bawa 1980; Givnish 1982) – although the hypothesis is controversial (Thomson & Barrett 1981; Charlesworth 1985). Either way, males and females of dioecious populations end up with phenotypes that enhance their own sex function, but that might not do so for the other, and might indeed be deleterious (Connallon & Clark 2014).

**Figure 6.**
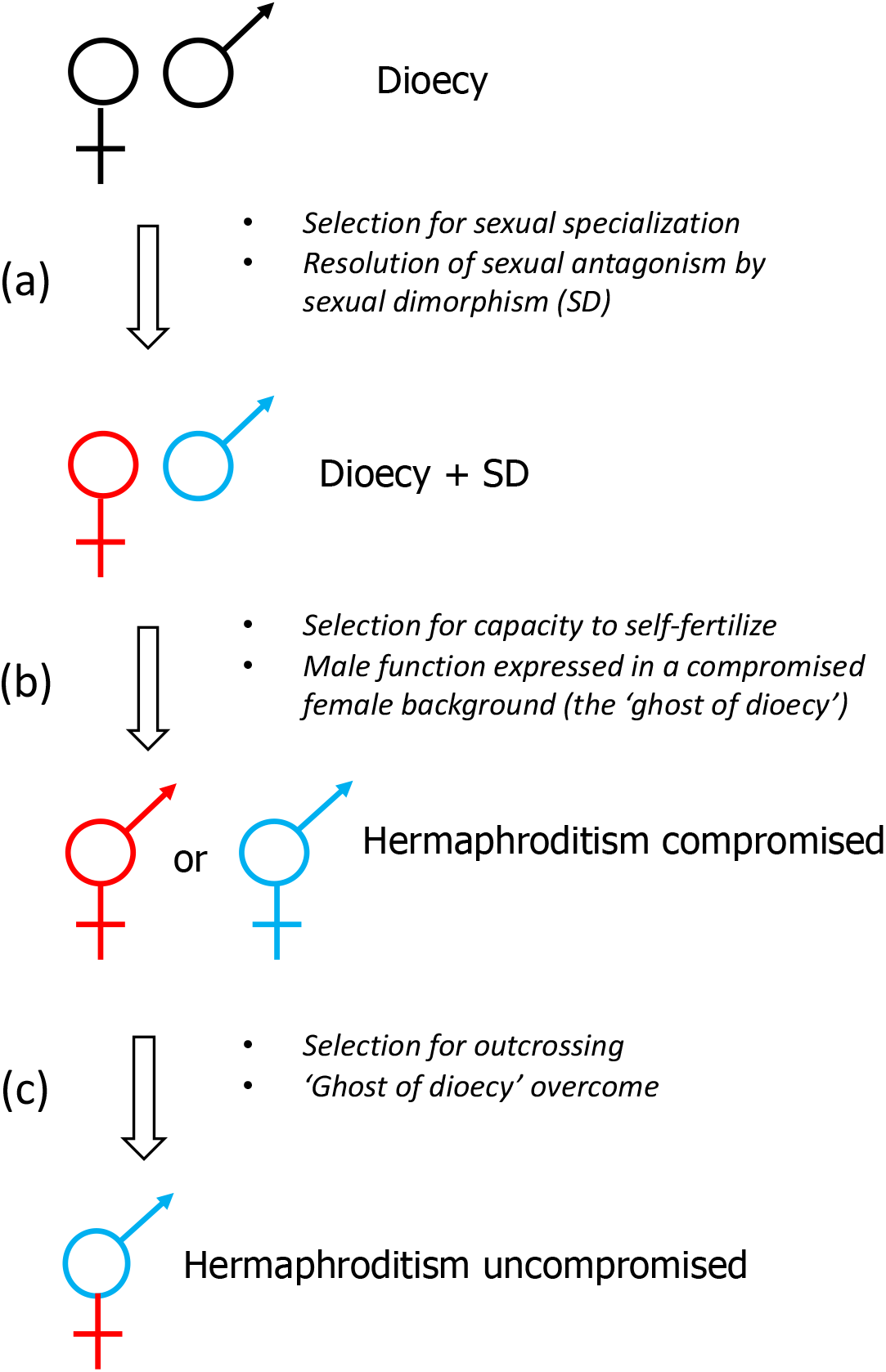
General evolutionary path envisaged for the breakdown of dioecy in flowering plants. (a) Over the course of its persistence, a dioecious population evolves secondary sexual dimorphism, with males and females expressing phenotypes that are optimized for their respective sexual functions. (b) Dioecy breaks down in response to selection for reproductive assurance and a capacity to self-fertilize by individuals with leaky sex expression. The hermaphrodites thus formed must express their newly acquired sex function in the context of a phenotype optimized for the other sex function. This compromised phenotype is what we refer to as the ‘ghost of dioecy’. (c) With time, selection for improved reproductive performance through both sexual functions molds the hermaphroditic strategy, finding phenotypes that improve one sexual function without, or with diminished, deleterious effects to the other.

The frequent transitions from dioecy to hermaphroditism in flowering plants (Kafer *et al*. 2014; Goldberg *et al*. 2017; Kafer *et al*. 2017) presumably begin with selection of males or females with a leaky sexual function, and the derived hermaphrodites (or monoecious individuals) should inherit the secondary sexual phenotype associated with their ancestral sexual function, male or female (Figure 6b). Our results suggest that this ancestral phenotype is antagonistic to the newly acquired function in monoecious *M. annua*, and similar reasoning may hold for transitions from dioecy to hermaphroditism more generally. For example, many dioecious lineages may revert to hermaphroditism via gynodioecy through the selection of leaky males, which are much more common than leaky females (Ehlers & Bataillon 2007; Cossard *et al*. 2018). In these lineages, hermaphrodites derived from leaky males will tend to express their acquired female function in the context of floral, physiological, life-history or defense traits that have been selected to optimize siring success, not seed or fruit production. With time, we expect selection to mold the hermaphroditic phenotypes in ways that overcome this ghost of dioecy, finding solutions that optimize both male and female functions and, for example, avoiding their interference (Figure 6c).

The above reasoning implies that sexual dimorphism should constrain the breakdown of dioecy when outcrossing is favored by establishing populations with a concave fitness set that become more resistant to the invasion of hermaphrodites. Indeed, once sexual dimorphism has evolved, it is difficult to imagine what could allow a transition from dioecy to hermaphroditism without an accompanying change in the mating system. In the case of *M. annua*, monoecy likely evolved from dioecy as a selfing mechanism in response to selection for reproductive assurance in sparse populations (Hesse & Pannell 2011a; Labouche *et al*. 2015), during episodes of range expansion (Pujol & Pannell 2008; Pujol *et al*. 2009), or during colonization of disturbed habitat in metapopulations (Pannell 2001; Pannell & Dorken 2006; Pannell *et al*. 2008), and mate limitation has been invoked for the breakdown of dioecy in other plant and animal lineages (Baker & Cox 1984; Charlesworth 1993; Maurice & Fleming 1995; Pannell 2002; Wolf & Takebayashi 2004; Ehlers & Bataillon 2007; Crossman & Charlesworth 2014). Our study emphasizes that such advantages must overcome the sometimes substantial benefits of sexual specialization enjoyed by males and females. Thus, while reproductive assurance may favor transitions from dioecy to hermaphroditism, sexual dimorphism (a legacy of selection under dioecy), should resist such transitions in outcrossing populations. We suggest that dioecy is thus most likely to break down when accompanied by a shift from outcrossing to selfing or mixed mating.

## Acknowledgements

We thank Nicolas Ruch for help with the array experiments, Laure Olazcuaga for help in the laboratory and field, Crispin Jordan, Paris Veltsos, Wen-Juan Ma, Marcos Méndez Iglesias, and two anonymous referees for comments on the manuscript, and the Swiss National Science Foundation for funding to JRP.

